# Evolved differences in microglial cell biology between surface and cave populations of *Astyanax mexicanus*

**DOI:** 10.64898/2026.04.11.717796

**Authors:** Emilio Mendez Scolari, Othniel K. Amanyi, Aakriti Rastogi, Erik R. Duboue, Alex C. Keene, Harini Iyer

## Abstract

Microglia govern multiple aspects of brain architecture and function by eliminating dying cells, stimulating neurogenesis, refining neural connections, and orchestrating immune responses. The Mexican tetra, *Astyanax mexicanus*, is a powerful model system for investigating the evolution of brain function, yet microglia have not been investigated in this system. *A. mexicanus* exists as surface-dwelling and cave morphotypes with prominent behavioral and physiological differences. Notably, these evolved behavioral and physiological changes in cavefish, including diminished immune response, sleep, circadian rhythms, and sensory processing, are directly linked to known microglial functions. These observations suggest that evolved differences in microglia may shape brain circuitry adaptations in cavefish. Here we develop an experimental toolbox to examine microglial specification, dynamics, and function in *A. mexicanus* to perform comparative analysis of microglial cell biology between the surface and cave morphotypes. We find that the cave populations show increased numbers of microglia over developmental time relative to their surface counterparts. Microglia in *Astyanax* rapidly expand in response to inflammatory cues, distinct from microglial responses in the related teleost, zebrafish. Furthermore, lysosomal compartments of microglia in the cave populations exhibit increased enhanced proteolytic activity and reduced pH relative to surface morphotypes. Together, our observations reveal evolved differences in microglial cell biology between surface and cave populations of *A. mexicanus* and provide a framework to uncover novel neuroimmune mechanisms underlying the remarkable adaptations of *A. mexicanus*.

## Introduction

Microglia, the brain’s resident sentinel immune cells, serve as first responders to perturbations in the central nervous system **(Prinz et al., 2019)**. Although derived from macrophages, microglia adopt highly specialized, brain-specific functions that extend beyond classical immune surveillance **(Cowan and Petri, 2018)**. During development, microglia support synaptic maturation **(Schafer et al., 2012)**, promote neurogenesis **(Shigemoto-Mogami et al., 2014)**, and eliminate excess or apoptotic cells **(Cunningham et al., 2013)**. In the adult brain, they continuously survey the parenchyma **(Nimmerjahn et al., 2005)**, modulate myelination **(Hagemeyer et al., 2017)**, mediate repair **(Bellver-Landete et al., 2019)**, and play a critical role in shaping normal behaviors **(Rogers et al., 2011; Zhan et al., 2014)**. The diverse and essential roles of microglia in central nervous system development and homeostasis suggest that microglia may contribute to the evolution and adaptation of brain circuits to niche-specific environments, thereby influencing animal physiology and behavior. Yet, the contributions of microglia to brain evolution remain largely unexplored in most evolutionary model systems.

The Mexican tetra, *Astyanax mexicanus*, has emerged as a powerful comparative model system for studying extreme trait evolution **(Jeffery et al., 2023; Moran et al., 2023; Oliva et al., 2022; Warren et al., 2024)**. This species exists as surface- and cave-dwelling morphotypes that share a common genetic background but experience profoundly different environments, resulting in the repeated evolution of distinct phenotypes. The cave morphs exhibit well-characterized traits such as albinism **(Protas et al., 2006)**, eye loss **(Choy et al., 2025; Shennard et al., 2025)**, and reduced sleep **(Duboue et al., 2011; Gallman et al., 2024; Jaggard et al., 2018; Keene and Duboue, 2018; Lloyd et al., 2025)**. Furthermore, differences between the morphotypes in metabolism **(Iwashita et al., 2023; Krishnan et al., 2022)**, sex-specific gene expression **(Webster et al., 2025)**, behavior **(Chin et al., 2018; Elipot et al., 2013; Iwashita and Yoshizawa, 2021; Yoshizawa et al., 2010)**, and aging **(Cobham et al., 2025)** have expanded the utility of this system for comparative cellular, physiological, and behavioral analyses. Together, the robust phenotypic differences between surface fish and cavefish provide an opportunity to identify the genetic and cellular basis of extreme trait evolution.

Importantly, microglial functions are intimately linked to multiple traits that differ between surface and cave morphotypes of *Astyanax*, including sleep **(Ma et al., 2024)**, aggression **(Takahashi, 2024)**, metabolism **(Bernier et al., 2020; Yang et al., 2021)**, aging **(Li et al., 2023; Norden and Godbout, 2013)**, and sex-specific behaviors **(Bordt et al., 2020; Thion et al., 2018b)**. However, microglial biology in the Mexican tetra remains poorly understood, limited by the availability of tools to interrogate these critical neuroimmune glia. A defining feature of microglial identity is that it is uniquely informed by developmental ontogeny and the brain environment, necessitating the study of microglia in vivo **(Bennett et al., 2018; Bohlen et al., 2017; Cronk et al., 2018; Gosselin et al., 2014)**.

Here we develop an experimental toolbox for assaying microglial function in *A. mexicanus*. Using live imaging to visualize microglia in vivo, we perform a developmental time course of the colonization of these essential neuroimmune cells in the brain. Our data show that as development proceeds, cave morphs exhibit significantly higher numbers of microglia than surface morphs. Notably, when challenged with inflammatory stress, microglia in *Astyanax* display robust proliferative responses. Furthermore, cavefish microglia show distinct lysosomal properties such as increased proteolytic enzyme abundance and reduced pH. Thus, our comparative analyses reveal important differences in brain-resident innate immune cells between *A. mexicanus* morphotypes. Collectively, our experiments establish methods to integrate existing behavioral and genomic resources in *Astyanax* with cell biological approaches to elucidate neuroimmune mechanisms underlying extreme trait evolution.

## Materials and Methods

### Astyanax mexicanus husbandry

Animal husbandry was carried out as previously described **(Kozol et al., 2023)**. Fish were housed in the Texas A&M core facilities at 23 ± 1°C constant water temperature throughout rearing. The animals were kept on a 14:10 hour light-dark cycle that remained constant throughout the animal’s lifetime. Light intensity was maintained between 25 and 40 lx for rearing. Adult fish were fed a diet of black worms to satiation twice daily at zeitgeber time (ZT) 2 and ZT12 (Aquatic Foods), and standard flake fish food during periods when fish were not being used for breeding (Tetramin Pro).

### Astyanax mexicanus and Danio rerio microscopy

To keep the protocols consistent across *D. rerio* and *A. mexicanus*, surface (Río Choy) and cave (Pachón) morphotype larvae were maintained in 100 mm Petri plates at a density of 50-60 larvae per dish at 23 °C. Similar to zebrafish, the *Astyanax* larvae were kept in embryo water with methylene blue and treated with 0.003% PTU starting at ∼1 day post fertilization (dpf) to inhibit pigmentation. Although cave morphs do not develop pigmentation, PTU treatment was applied uniformly to ensure consistent conditions across all groups. Zebrafish AB wild-type embryos were raised at 28.5 °C in embryo water with methylene blue and treated with 0.003% PTU starting at 20-24 hours post fertilization (hpf) to inhibit pigmentation. All the animal protocols were approved by Rice University Institutional Animal Care and Use Committee.

### Neutral red assay

A 2.5 mg/ml neutral red (Sigma, N7005) stock solution was prepared by dissolving the dye in ultrapure distilled water and filtering the solution through a 0.45 µm syringe filter. For the staining, larvae were incubated for 2 hours in embryo water containing 5 µg/ml of neutral red in embryo water containing PTU. Following two rinses with embryo water to remove any remaining dye, larvae were washed overnight in embryo water with PTU. After approximately 18 to 20 hours of incubation, larvae were anesthetized with 0.016% MS222 (Tricaine) and immediately mounted in 1.5 % low-melting-point agarose. Images were acquired using a Zeiss Stemi 508 stereomicroscope. Area of the midbrain was measured using Fiji software (ImageJ, NIH, USA). Statistical analyses and graph generation were performed using GraphPad Prism (GraphPad Software, San Diego, CA, USA).

In the time course experiment, for the 1 and 2 dpf time points, zebrafish embryos were dechorionated by incubation for 5 minutes in 1 mg/ml Pronase (Roche, 10 165 921 001), dissolved in embryo water containing PTU. Dechorionated embryos were washed several times with embryo water and allowed to recover in embryo water with PTU for at least 4 hours prior to neutral red treatment. Cavefish and surface fish embryos were naturally dechorionated before 24 hpf.

### Tail injury

To induce tail injury, 4 dpf larvae were anesthetized and placed on a flat surface. The distal tip of the caudal fin was transected using a new scalpel. Incisions were performed to ensure a similar injury size across animals while avoiding damage to the spinal cord. At 4 hours post-injury, the neutral red assay was performed to stain macrophages. Following overnight wash and incubation (5 dpf), larvae were anesthetized with 0.016% MS222 (Tricaine) and mounted in 1.5% low-melting-point agarose to acquire images of the injured tail using a Zeiss Stemi 508 stereomicroscope. The number of neutral red-positive macrophages was quantified, and statistical analyses were performed using GraphPad Prism (GraphPad Software, San Diego, CA, USA).

### LysoTracker Red and LysoSensor Green assay

LysoTracker Red DND-99 (1 mM, Thermo Fisher Scientific, L7528) and LysoSensor Green DND-189 (1 mM, Thermo Fisher Scientific, L7535) were diluted in embryo water with PTU to get 1 µM working solution (1:1000). Larvae at 4-6 dpf were incubated with the corresponding dye for 45 minutes on a shaker in dark. Following staining, larvae were washed three times in embryo water containing PTU, 15 minutes per wash, on agitation and in dark. Immediately after washing, larvae were anesthetized with 0.016% MS222 (Tricaine), mounted in 1.5% low-melting-point agarose, and imaged using a Zeiss LSM 900 confocal microscope. The fluorescence intensity of LysoTracker Red and LysoSensor Green puncta was measured using Fiji software (ImageJ, NIH, USA). Statistical analyses and graph generation were performed using GraphPad Prism (GraphPad Software, San Diego, CA, USA).

### Brain injections

For brain microinjection, anesthetized larvae were aligned in a 1.5% grooved agarose mold, with the dorsal side facing upward to allow access to the brain. One nanoliter of poly(I:C) (Tocris Bioscience, 4287; 1 mg/ml in 1X PBS), zymosan (Sigma-Aldrich, Z4250; 10 mg/ml in 1X PBS, boiled to solubilize), Magic Red Cathepsin (Immunochemistry Technologies, 937; vial 6133 suspended in 50 µL of DMSO and diluted 1:1 in 1X PBS before use), *E. coli* Texas Red (Thermo Fisher Scientific, E2863; 10 mg/ml in PBS), and Amyloid-β (1-42) HiLyte-Fluor 488 (Anaspec, AS-60479; 1 mg/ml in 1X PBS with 1% NH_4O_H) were injected into one hemisphere of the midbrain. A small volume of phenol red was added to each injection mix to aid with visualization during the injection. Amyloid-β (1-42) HiLyte Fluor-488 was injected at 4 dpf, and imaging was performed at 5 and 6 dpf (1 and 2 dpi, respectively). Poly(I:C) and zymosan were injected at 4 and 5 dpf, respectively. Magic Red cathepsin, and *E. coli* Texas Red were injected at 6 dpf. The injected animals were allowed to recover in embryo water with PTU for 2 to 4 hours prior to additional experiments or imaging.

### Immunohistochemistry

All larvae were fixed at 4 and 5 dpf in 4% PFA and 1% DMSO overnight. The different permeabilization and post-fixation conditions are specified in Table 1. After a single wash in 1× PBS, larvae were incubated for 2 hours in blocking solution (1% DMSO, 1% heat-inactivated donkey serum, 1% bovine serum albumin, and 0.7% Triton X-100 in 1× PBS) on a nutator. Larvae were then incubated overnight at 4 °C on a nutator with anti-Lcp1 (GeneTex, GTX124420) and anti-Mfap4 (GeneTex, GTX132692) antibodies (1:300 dilution in blocking solution). After six washes of 10 minutes each in 1× PBS containing 0.8% Triton X-100, larvae were incubated for 2 hours at room temperature on a nutator in blocking solution containing Alexa Fluor 488–conjugated donkey anti-rabbit IgG (1:500; Jackson ImmunoResearch, 711-545-152). Following six additional 10-minute washes in 1× PBS containing 0.8% Triton X-100, larvae were mounted for confocal imaging.

**Table 1.** Optimization of IHC permeabilization in fixed *Astyanax*. Comparative analysis of Lcp1 and Mfap4 signal detection across immunohistochemistry conditions.

| Experimental treatment |  |  |  | Microglia signal |  | Macrophage signal |  |
| --- | --- | --- | --- | --- | --- | --- | --- |
| Fix | Permebealization | Permeabilization time | Post-fixation | Lcp1 | Mfap4 | Lcp1 | Mfap4 |
| Fix 4% PFA, 1% DMSO | None | N/A | No | - | + | +++ | ++ |
|  | Collagenase (2 mg/ml) | 45 min | No | Not tested | + | Not tested | +++ |
|  | Proteinase K (80 µg/ml) | 20 min | No | - | - | - | - |
|  | Proteinase K (40 µg/ml) | 20 min | No | - | - | - | - |
|  | Proteinase K (20 µg/ml) | 45 min | No | - | +++ | + | - |
|  | Proteinase K (80 µg/ml) | 20 min | Yes | - | - | ++ | ++ |
|  | Proteinase K (40 µg/ml) | 20 min | Yes | - | + | + | + |
|  | Proteinase K (20 µg/ml) | 60 min | Yes | + | ++ | +++ | ++ |

Fluorescence signal was qualitatively scored as - (no signal), + (low), ++ (moderate), +++ (strong)

## Results

### Neutral red labels microglia and macrophages in *Astyanax mexicanus*

To define a protocol that labels microglia in *A. mexicanus* surface and cave populations, we performed neutral red staining across multiple stages of early development **(Fig. 1)**. Neutral red is a vital dye that accumulates in acidic compartments of the cell and is widely used for live imaging of microglia in zebrafish in the midbrain or optic tectum **(Herbomel et al., 1999; Herbomel et al., 2001)**. We therefore adapted the neutral red labeling protocol **(Iyer et al., 2022; Iyer and Talbot, 2024)** for use in *A. mexicanus*. As previously described, in zebrafish, the number of embryonic macrophages colonizing the midbrain increased steadily over developmental time, with the most pronounced increase occurring between 2-3 days post-fertilization (dpf) **(Iyer et al., 2022) (Fig. 1A, 1D, 1E)**. Similarly, both surface and cave morphs of *A. mexicanus* showed a progressive increase in the number of neutral red-labeled microglia in the midbrain over development **(Fig. 1B-1D)**. At early stages, surface fish exhibited higher numbers of neutral red-labeled microglia; however, by 5 dpf, the cave morph displayed significantly greater numbers of microglia relative to the surface morph **(Fig. 1B-1D)**. Therefore, neutral red effectively labels microglia in *A. mexicanus* and identifies differences between surface fish and cavefish.

**Figure 1.**
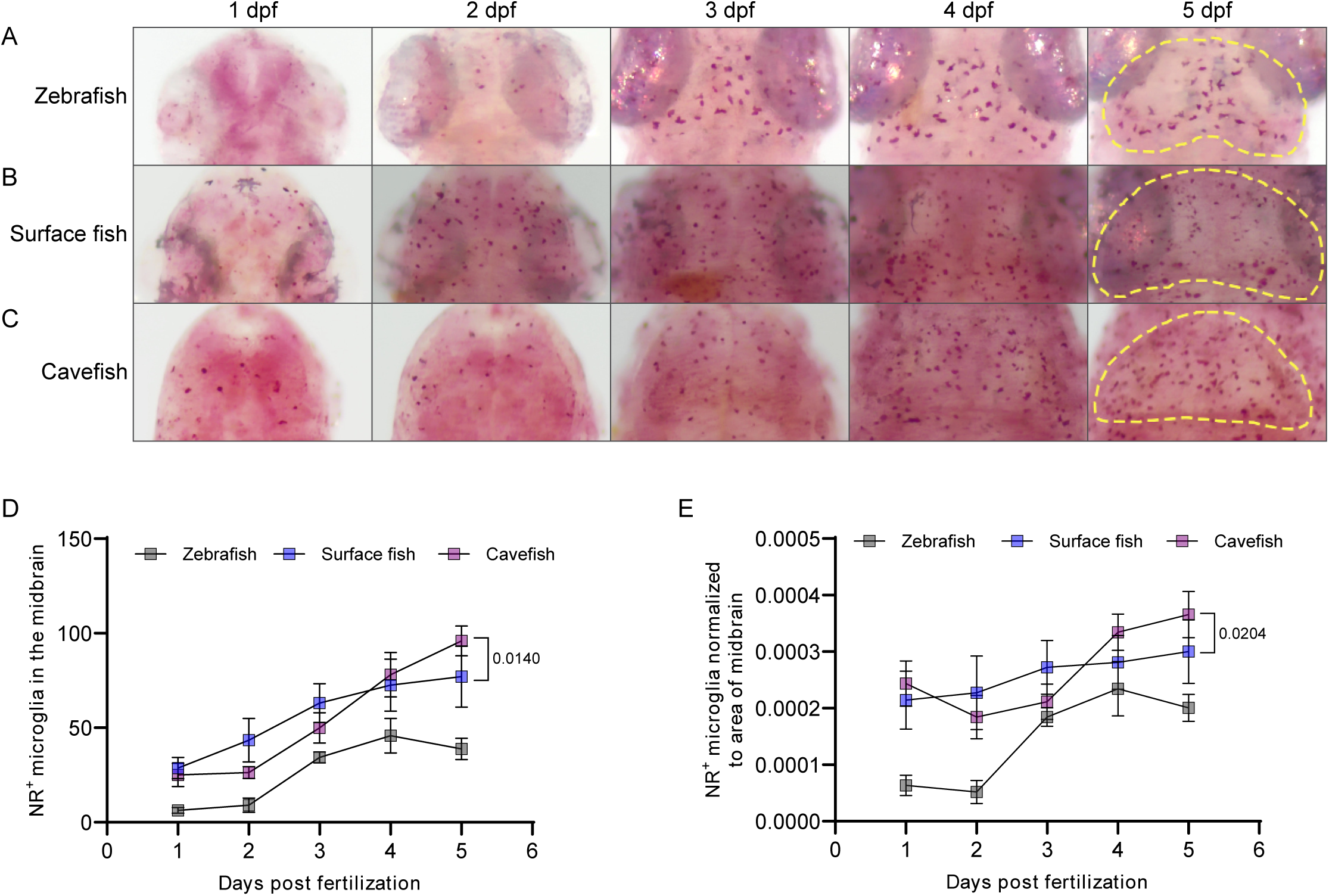
Neutral red labels microglia and macrophages in *Astyanax*. Neutral red labeling was performed across developmental time points in **(A)** Zebrafish **(B)** *A. mexicanus* surface morphs, and **(C)** *A. mexicanus* cave morphs. **(D)** Raw counts of neutral red-labeled microglia in the midbrain. **(E)** Microglial density calculated by normalizing raw microglia numbers to midbrain surface area in *D. rerio* and *A. mexicanus*. Each data point in panels **D** and **E** represents mean ± SEM from individual larva. Statistical analysis was performed using an unpaired *t*-test.

Because brain volume differs regionally as well as over development between surface and cave morphotypes, we quantified not only the raw numbers of neutral red-labeled cells in this region, but also microglial density (neutral red^+^ cell number normalized to midbrain area) across developmental stages **(Fig. 1E)**. Delineation of the midbrain is challenging at 1 dpf in both *Danio rerio* and *Astyanax mexicanus*; however, beginning at approximately 2.5 dpf, a distinct bean-shaped structure corresponding to the optic tectum becomes apparent **(Fig. 1A-1C)**. Consistent with the raw neutral red cell counts, microglial density increased over developmental time, and this increase in microglial density became significant by 5 dpf **(Fig. 1B, 1C, 1E)**.

Previous studies have reported that cave populations possess fewer cells of the myeloid lineage than their surface counterparts **(Peuss et al., 2020)**. To determine whether the increased microglial numbers we observed in cave morphotypes reflected broader differences in macrophage populations, we used neutral red labeling to examine peripheral macrophages in the tail region of surface and cave morphs. Consistent with prior reports, cave morphs exhibited markedly fewer macrophages in peripheral tissues compared to surface fish **(Fig. S1A, arrows**). To examine whether these differences in macrophage numbers led to altered functional responses, we performed tail fin injury of the larvae followed by neutral red labeling. Injury induced an increase in macrophage numbers in the tails of cavefish larvae **(Fig. S1B)**, but macrophages in both surface and cave morphs showed comparable recruitment to the injury site **(Fig. S1B, S1C)**, indicating that peripheral macrophage injury responses are largely similar between the surface and cave forms of *Astyanax* despite reduced macrophage numbers in cavefish in homeostasis. Collectively, these experiments demonstrate that microglial and macrophage populations in both surface and cave morphs of *A. mexicanus* can be labeled with neutral red and traced over developmental time.

### Neutral red labeling for tracking microglial activation in response to inflammatory stimuli

As the primary immune cells of the central nervous system, microglia are the first responders to brain perturbations **(Thion et al., 2018a)**. We therefore investigated microglial responses to inflammatory stimuli in *A. mexicanus* by modifying zebrafish microinjection techniques to deliver debris or pharmacological agents into tissues or circulation **(Rosen et al., 2009)**. Larvae were positioned in custom agarose molds, oriented with the dorsal surface facing the injection needle, and maintained in a small volume of water. A defined volume of inflammatory molecules was injected directly into the midbrain parenchyma **(Fig. 2A)**.

**Figure 2.**
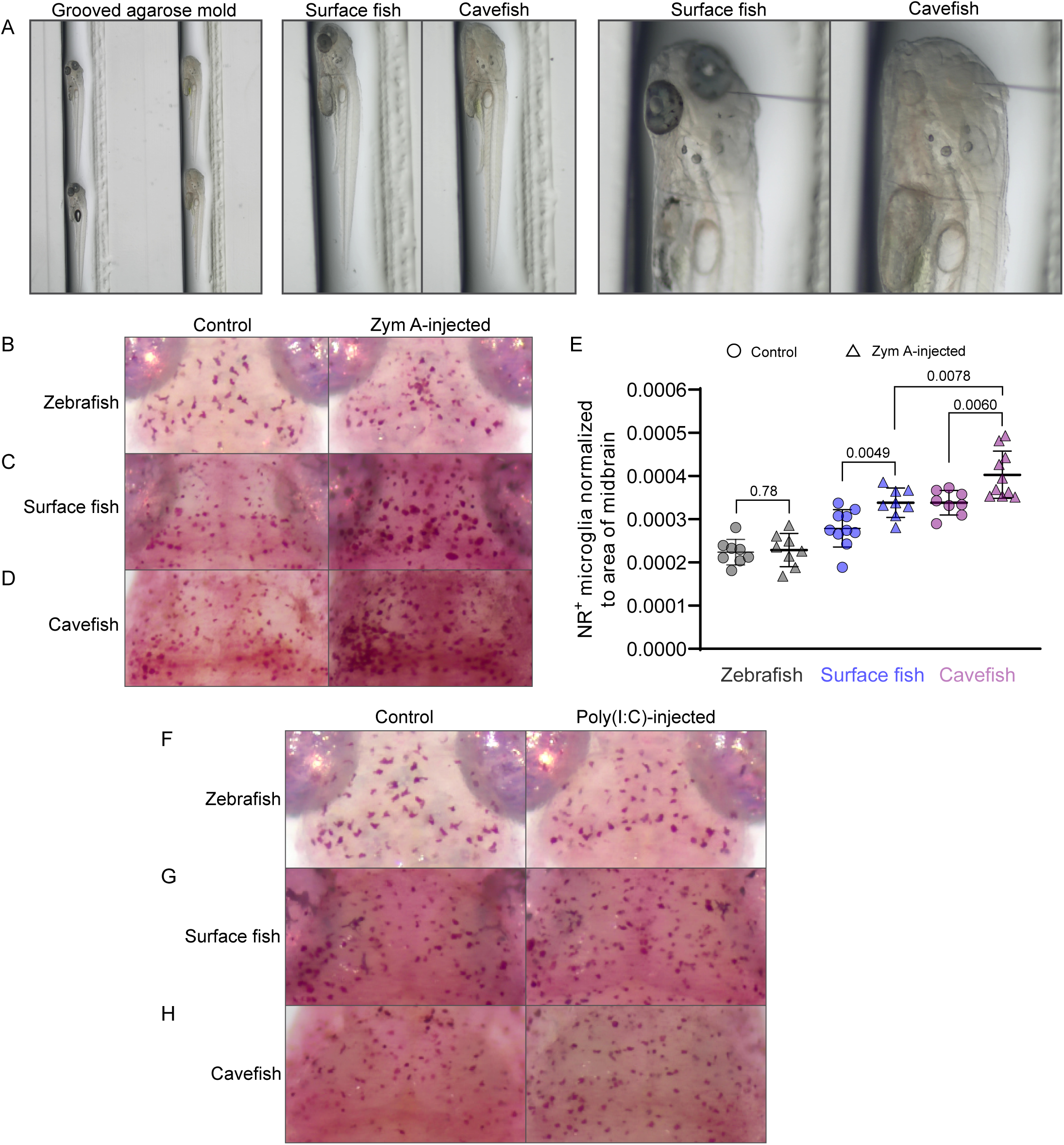
Distinct microglial responses to inflammatory stimuli in *Astyanax*. **(A)** Schematic of the microinjection setup. **(B–D)** Zymosan A injection into the brain parenchyma followed by neutral red staining in **(B)** Zebrafish, **(C)** *A. mexicanus* surface morphs, and **(D)** *A. mexicanus* cave morphs. **(E)** Quantification of microglial density in the midbrain following Zym A injection. Each data point represents a single larva. Graph shows mean ± SD. Statistical analysis was performed using an unpaired *t*-test. **(F-H)** Poly(I:C) injections in zebrafish, surface fish, and cavefish larvae followed by neutral red labeling.

Zymosan A is a potent immune stimulant derived from the yeast cell wall and is widely used to induce sterile inflammation. In zebrafish, macrophages and microglia rapidly respond to zymosan A (Zym A) injection by adopting an amoeboid morphology, a hallmark of microglial activation, and migrating to engulf zymosan particles **(Iyer et al., 2022)**. We therefore used Zym A injection to monitor microglial responses to inflammatory cues in *Astyanax*. Because the injection procedure itself is perceived as an injury signal by microglia, uninjected animals were used as controls. In both *D. rerio* and *A. mexicanus*, microglia exhibited robust responses to Zym A injection **(Fig. 2B-2D)**. In agreement with previous findings, zebrafish microglia transitioned from a ramified to a highly amoeboid morphology, consistent with activation **(Fig. 2B)**. In surface and cave morphs of *Astyanax*, microglia also displayed changes in cell shape following Zym A injection, although baseline microglial morphology in uninjected animals was less ramified than in zebrafish **(Fig. 2C, 2D)**.

A striking difference between *D. rerio* and *A. mexicanus* surface and cave populations was observed in microglial abundance following Zym A injection. Both surface and cave morphs of *Astyanax* showed a significant increase in microglial numbers, whereas zebrafish did not **(Fig. 2B-2E)**. To assess the extent to which this response was specific to Zym A or represented a more general inflammatory adaptation in *A. mexicanus,* we injected poly(I:C), a synthetic double-stranded RNA analog that mimics viral infection and activates innate immune signaling, into the brain parenchyma. Similar to Zym A treatment, poly(I:C) injection resulted in expanded microglia in both surface and cave morphs of *A. mexicanus* **(Fig. 2G, 2H)** relative to uninjected controls and minimal changes in microglia quantity was observed in zebrafish **(Fig. 2F)**. Together, these data demonstrate that microglia in *A. mexicanus* mount robust activation in response to inflammatory stimuli, but responses in both surface and cave morphs are distinct from those observed in zebrafish.

### Labeling microglia and macrophages using immunohistochemistry

Although live imaging is optimal for visualizing microglial and macrophage functions such as surveillance, chemotaxis, and debris clearance, many experiments, including assays of cell proliferation (e.g., EdU labeling) or cell death (e.g., TUNEL), require fixed tissue. We therefore optimized immunohistochemistry protocols **(Table 1)** to label microglia and macrophages in fixed *A. mexicanus* larvae. We used antibodies widely employed in zebrafish, anti-Lcp1 **(Jin et al., 2009; Lopez-Munoz et al., 2018)** and anti-Mfap4 **(Campbell et al., 2024; John et al., 2025)**, to label both microglia and macrophages.

Following fixation with 4% PFA, we tested both collagenase and proteinase K treatments to permeabilize larvae. In general, proteinase K yielded a higher signal-to-noise ratio compared to collagenase. We further tested a range of proteinase K concentrations, as well as inclusion of a post-fixation step, and observed the highest signal at 20 µg/ml proteinase K **(Table 1)**. As anticipated, macrophages in the tail region were readily detected using both anti-Lcp1 and anti-Mfap4 **(Fig. 3A, 3B),** likely due to efficient permeabilization of this thin tissue. Microglia were more difficult to detect overall, likely due to rapid skull development and the deeper positioning of microglia, and we observed more robust labeling with both anti-Lcp1 and anti-Mfap4 in cavefish relative to their surface counterparts **(Fig. 3C).**

**Figure 3.**
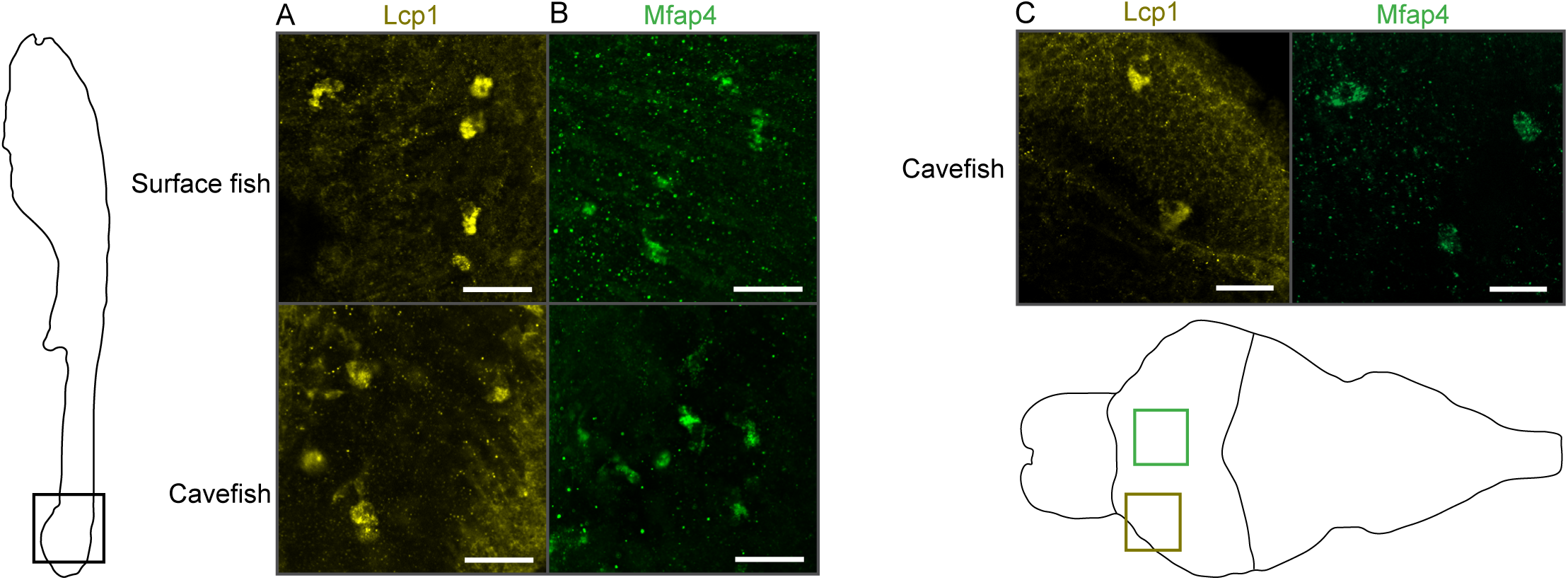
Examining macrophages and microglia using immunohistochemistry in *Astyanax*. Anti-Lcp1 and anti-Mfap4 antibody labeling of **(A, B)** macrophages and **(C)** microglia. Scale bars: 25 µm.

### Distinct lysosomal compartment features in cavefish microglia

The neutral red assay is a convenient method for evaluating gross changes in numbers or distribution of microglia and macrophages. However, analysis of debris engulfment and lysosomal proteolytic capacity requires fluorescent probes that allow subcellular resolution and tracking. Several commercially available dyes enable high-resolution imaging of lysosomal compartments within microglia and have been widely used in zebrafish **(Iyer et al., 2022)**.

LysoTracker Red and LysoSensor Green label acidic lysosomal compartments in zebrafish microglia **(Iyer et al., 2022) (Fig. 4A)**. We therefore optimized concentrations and treatment protocols for these dyes in both surface and cave morphs of *A. mexicanus*. Live imaging of single coronal sections revealed robust labeling of microglia with both dyes in surface and cave populations **(Fig. 4B, 4C)**. Notably, fluorescence intensity of both LysoTracker and LysoSensor was greater in cavefish relative to surface fish **(Fig. 4D, 4E)**, indicating either expanded lysosomal compartments or increased lysosomal acidity in the microglia of the cave morph. Although our confocal imaging was limited to single optical planes, LysoTracker and LysoSensor labeling corroborated our neutral red observations, revealing increased numbers of microglia in cave morphs compared to surface morphs **(Fig. 1)**.

**Figure 4.**
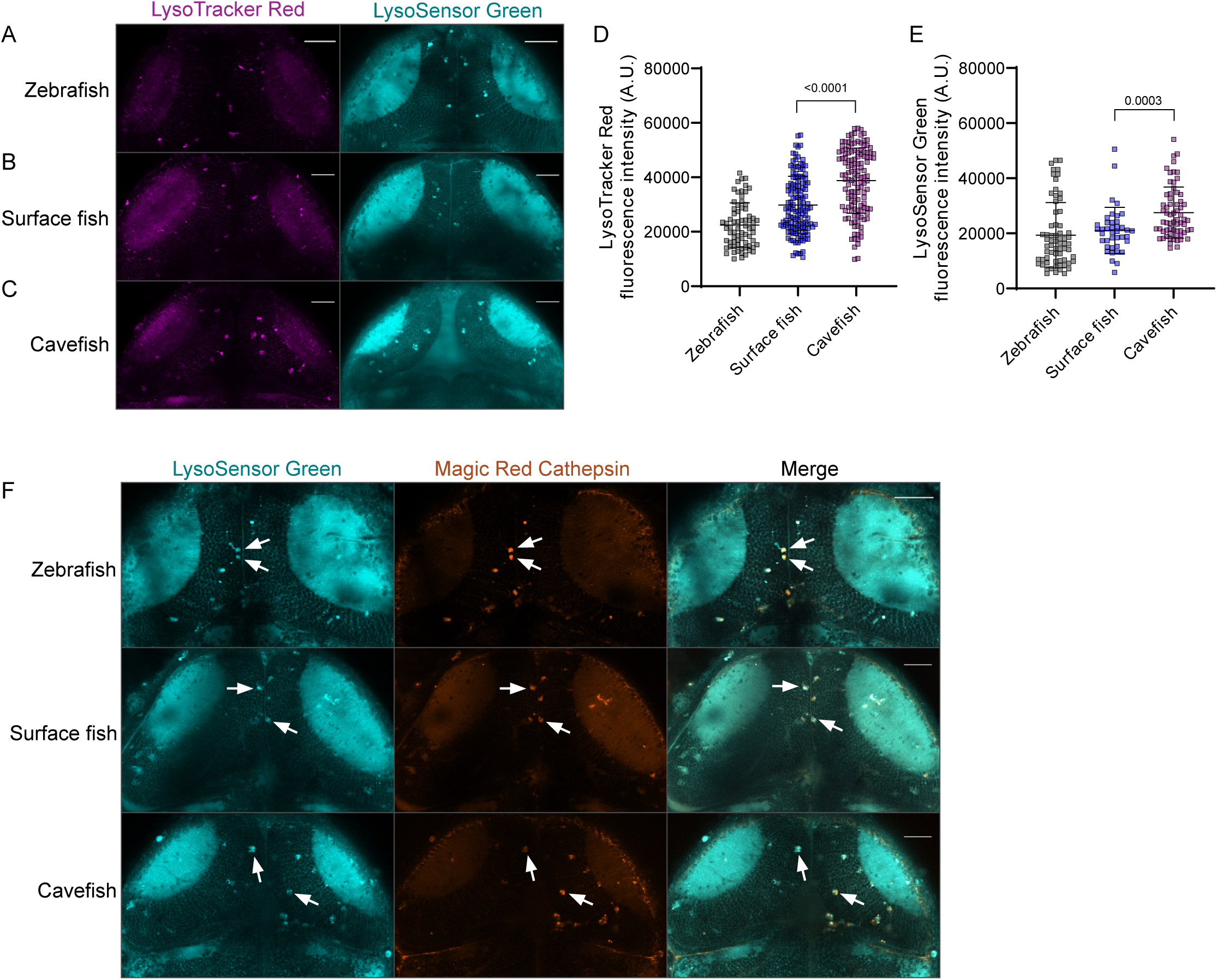
Visualizing lysosomal compartments in microglia in *Astyanax*. (A–C) *D. rerio*, *A. mexicanus* surface morph, and *A. mexicanus* cave morph larvae were treated with LysoTracker Red and LysoSensor Green to visualize lysosomal compartments in microglia using live imaging. Scale bars: 50 µm. **(D-E)** Quantification of fluorescence intensity of: **(D)** LysoTracker Red and **(E)** LysoSensor Green puncta. Each data point represents a intensity from a single puncta. Graphs show mean ± SD. Statistical analysis was performed using an unpaired *t*-test. **(E)** Dual labeling with LysoSensor Green and Magic Red Cathepsin to assess proteolytic compartments within microglial lysosomes. Arrows indicate colocalization of LysoSensor Green with Magic Red Cathepsin within microglia. Scale bars: 50 µm.

We next examined lysosomal proteolytic enzyme abundance in microglia using Magic Red Cathepsin, a fluorogenic substrate that is normally quenched but becomes fluorescent upon cleavage by active lysosomal proteases **(Iyer et al., 2022)**. Unlike neutral red, LysoTracker Red, or LysoSensor Green, Magic Red Cathepsin is not efficiently taken up by cells through soaking. We therefore injected Magic Red Cathepsin directly into the brain parenchyma of surface and cave larvae, followed by LysoSensor Green labeling to detect colocalization **(Fig. 4F)**. Robust Magic Red Cathepsin signal was observed in the midbrain of both surface and cave morphs and overlapped extensively with LysoSensor-positive compartments **(Fig. 4F)**. Consistent with our earlier observations, cave morphs exhibited a greater number of LysoSensor-and Magic Red Cathepsin-positive puncta compared to surface morphs **(Fig. 4F)**. Collectively, these experiments indicate that microglia in cave morphs possess expanded and proteolytically active lysosomal compartments relative to those in surface morphs.

### Microglial engulfment of microbial and protein debris in *Astyanax*

Microglia act as the primary phagocytic cells of the central nervous system and are responsible for sensing, engulfing, and degrading a wide range of exogenous and endogenous substrates. To more directly interrogate the functional phagocytic capacity in *A. mexicanus* microglia, we challenged larvae with two biologically distinct substrates: fluorescently labeled *Escherichia coli*, a canonical bacterial target of innate immune cells, and amyloid-β, a disease-relevant aggregate associated with neurodegenerative pathology. *E. coli* provides a robust readout of classical immune phagocytosis, whereas amyloid-β scavenging reflects microglial handling of endogenous protein aggregates and has been widely used to probe microglial function in zebrafish models of neurodegeneration **(Bhattarai et al., 2017; Bhattarai et al., 2022; Iyer and Talbot, 2024)**.

To visualize the localization of injected particles within lysosomal compartments, we paired injection of these exogenous debris molecules with LysoSensor and LysoTracker staining. Microglia in both surface and cave morphs robustly engulfed *E. coli* **(Fig. S2)** and amyloid-β **(Fig. 5A, 5C)**, with no detectable differences in uptake or intracellular signal intensity. These data therefore suggest that microglia in both surface and cave morphotypes display comparable capacity to engulf microbial and protein substrates. When we compared amyloid-β intensity between 1 and 2 days post-injection (dpi), we observed a significant reduction in engulfed amyloid-β in both surface and cave larvae, as indicated by the intensity of amyloid-β⁺ puncta colocalized with LysoTracker Red **(Fig. 5B, 5D)**, indicating clearance of the amyloid-β debris over time. We observed a more significant decrease in amyloid-β intensity in surface larvae relative to the cave morphotype. To observe the uptake of amyloid-β particles by migrating LysoTracker Red puncta, we performed timelapse imaging of amyloid-β-injected animals co-stained with LysoTracker Red **(Video S1).** Our timelapse movies show the uptake of amyloid-β particles by migrating LysoTracker Red puncta (yellow arrows, **Fig. 5E**). Together, these observations demonstrate that microglial chemotaxis, phagocytosis, and cleance can be effectively tracked in both surface and cave morphotypes of *A. mexicanus* using LysoTracker Red. These approaches provide the future basis of examining clearance across different genetic backgrounds and strains of *A. mexicanus*.

**Figure 5.**
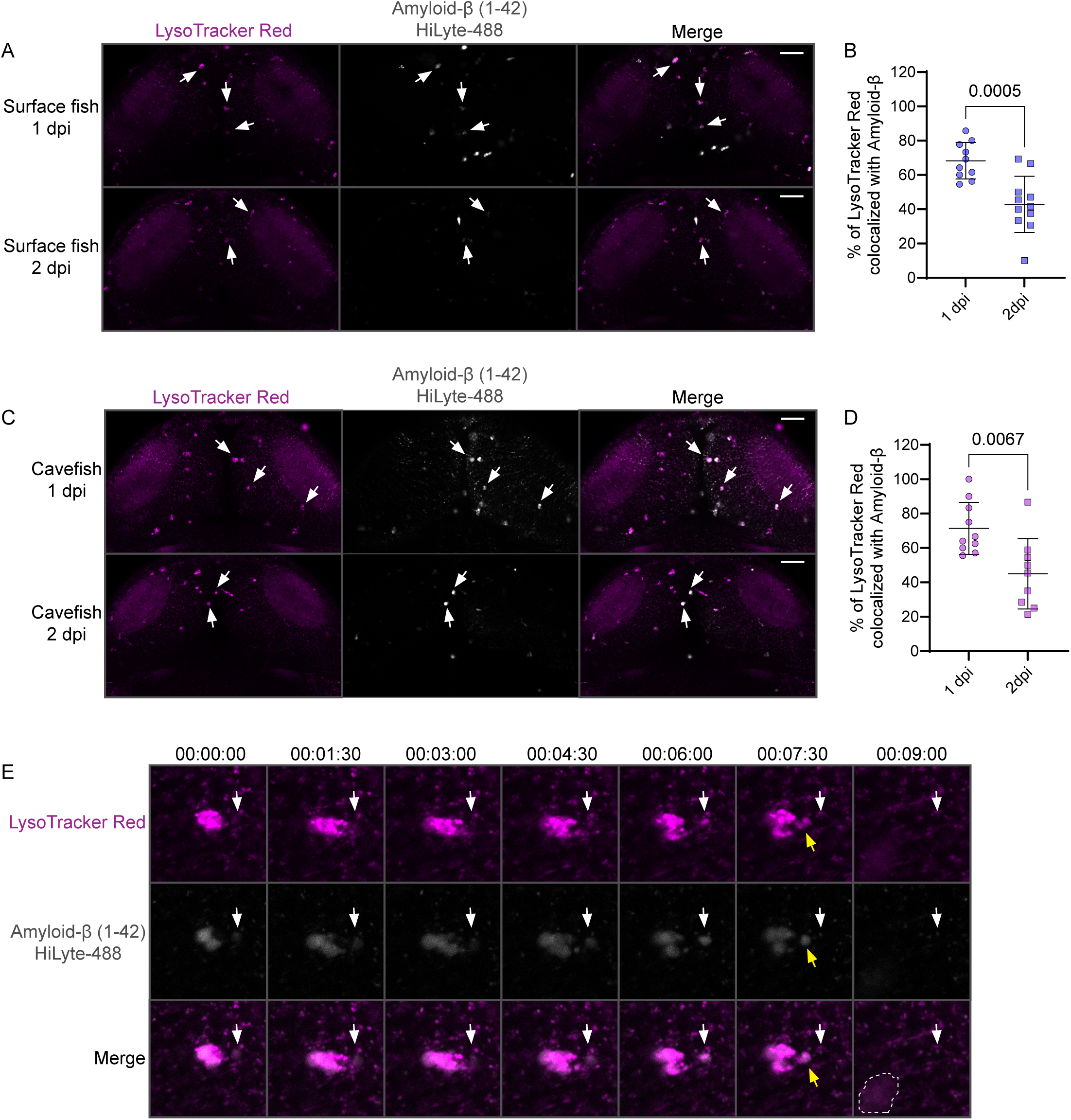
Exogenous debris engulfment in *Astyanax*. Injection of amyloid-β (1-42) HiLyte Fluor 488 into the brain parenchyma of **(A, B)** surface and **(C, D)** cave morphotypes followed by quantification of intensity. Scale bars: 50 µm. **(E)** Frames of timelapse imaging experiment. White arrows denote amyloid-β particles that are targeted by microglia for engulfment; yellow arrows show engulfed amyloid-β debris. The microglial cell, with the amyloid-β debris, moves out of frame in the last panel and its final location in the movie is indicated by a dotted shape.

## Discussion

Many key insights into microglial development, colonization, and function have been derived from teleost models, particularly zebrafish, which are highly amenable to live imaging and genetic manipulation **(Casano et al., 2016; Wu et al., 2018; Xu et al., 2016; Xu et al., 2015)**. The zebrafish community-developed tools have been instrumental in advancing neuroimmune research and provide opportunities for extending microglial studies to additional teleost systems **(Lam, 2022; Mazzolini et al., 2019; Peri and Nusslein-Volhard, 2008; Vanhunsel et al., 2021; Yasuda et al., 2015)**. Here, we adapt and optimize tools for studying microglia in the Mexican Tetra and identify differences in microglial properties between surface and cave morphotypes, as well as between *D. rerio* and *A. mexicanus*. Notably, we observed a marked increase in microglial numbers in cave larvae compared to their surface counterparts. In zebrafish, the best-established extracellular cues guiding embryonic microglia migration and brain colonization include apoptotic neuronal debris **(Casano et al., 2016; Xu et al., 2016)** and signaling through the Csf1r-Il34 pathway **(Wu et al., 2018)**. One possible explanation for the increased microglial numbers observed in cave morphs is altered regulation of these signaling pathways between surface and cave populations. Beyond serving as a pH-sensitive dye, neutral red also reports efferocytosis activity, as microglia must actively endocytose the dye. Although the increased number of microglia observed in cave morphs could reflect altered lysosomal pH or enhanced efferocytic capacity, cave-specific adaptations may also contribute to or arise from these differences in microglial abundance.

We also observed a clear difference in the microglial response to inflammatory stimuli between *D. rerio* and *A. mexicanus*. Injection of zymosan A induced a robust increase in microglial numbers in *A. mexicanus* larvae, whereas zebrafish microglia did not show a comparable expansion in response to the same stimulus. This divergence could reflect species-specific differences in how microglial populations are regulated during inflammation. In *A. mexicanus*, the increase in microglial numbers may arise from local proliferation, enhanced recruitment of peripheral macrophages, or prolonged retention/survival of activated microglia at the site of injury and inflammation. In contrast, zebrafish microglia may rely more on changes in activation state, such as altered morphology, migration, or phagocytic activity, rather than cell number expansion. These differences suggest that the balance between microglial proliferation and functional activation is tuned differently across species, potentially reflecting adaptation to distinct environmental or physiological pressures.

One of our most striking findings was the difference in lysosomal properties of microglia between cave and surface morphotypes. Multiple mechanisms could underlie this observation. Cave environments impose prolonged nutrient limitation and reduced sensory input **(Fernandes et al., 2022; Hamm and Gross, 2025; Powers et al., 2020)**. Microglia may therefore compensate by adopting a more catabolic state with increased lysosomal abundance, acidification, or protease activity to maximize recycling of cellular debris and extracellular material. Similarly, a shift toward glycolytic or lipid-dependent metabolism, a well-known cave adaptation **(Iwashita et al., 2023)**, could drive changes in lysosomal dynamics and signaling. Another interesting possibility is that neural circuit refinement or maintenance likely differ in cavefish due to reduced visual system input **(Yoshizawa et al., 2012)**. Microglia, particularly those in the optic tectum, may engage more heavily in synaptic pruning or debris clearance, necessitating enhanced lysosomal function to process engulfed material efficiently. Lastly, a constitutively “primed” microglial state in cavefish could lead to expanded or more active lysosomal compartments because lysosomes are central to antigen processing, phagolysosomal maturation, and inflammatory signaling.

Evolution of immune function is critical for adaptation to distinct ecological niches and social environments **(Gerardo et al., 2020; Mayer et al., 2016)**. The diverse roles of microglia in neuroimmune responses raise the possibility that they contribute to brain evolution, yet microglial biology has not been systematically examined in most evolutionary model systems. Tetra fish provide a powerful system to study ecological evolution and niche exploitation **(McGaugh et al., 2020)**, offering an opportunity to investigate how evolved changes in brain function and occupation of novel environments are associated with microglia. Here we have developed a comparative framework for connecting microglial states with adaptive behavioral evolution and neural circuit remodeling between surface and Pachón cave population. These studies can be readily extended to additional cave populations, providing opportunities to examine how variation in microglia function is associated with other physiological processes and evolutionary history.

In summary, our experiments establish tools and strategies for studying microglia in *A. mexicanus,* and we identify important differences in these critical neuroimmune cells between surface and cave morphotypes. At the same time, our findings open the field to several compelling research directions. What is the contribution of increased microglial abundance and altered lysosomal function to cave-specific adaptations? Why do cave morphotypes exhibit elevated microglial numbers in the brain while limiting innate immune populations in peripheral tissues? At the molecular level, which signaling pathways drive increased microglial abundance in cave populations, and what underlies the distinct microglial responses observed between *D. rerio* and *A. mexicanus*? Most importantly, how do these differences in microglial dynamics shape cavefish physiology, behavior, and adaptation to extreme environments? Together with future transgenic tools and the existing genomic and behavioral toolkit for cavefish, our experiments provide a framework to uncover cellular mechanisms underlying the remarkable adaptations of *A. mexicanus*.

## Legends

**Supplementary figure 1. Neutral red labels to visualize macrophage injury response in *Astyanax.* (A)** Neutral red labeling of peripheral macrophages in *A. mexicanus* surface and cave morphs. Arrows show macrophages in the caudal vein. **(B)** Tail fin injury assay in *Astyanax*. Dotted boxes indicate macrophages accumulation at the wound site. **(C)** Quantification of macrophage numbers at the injury site in surface and cave morphs. Each data point represents macrophage counts from a single larva. Graphs show mean ± SD. Statistical analysis was performed using an unpaired *t*-test.

**Supplementary figure 2. Bacterial debris engulfment in *Astyanax*.** *D. rerio* and *A. mexicanus* larvae were injected with *E. coli* Texas Red particles, followed by LysoSensor Green labeling to assess localization of *E. coli* within lysosomes. Arrows show colocalization of LysoSensor Green with *E. coli* Texas Red. Scale bars: 50 µm.

**Video S1. Time-lapse imaging of amyloid-β engulfment in *Astyanax*.** Amyloid-β (1–42) HiLyte Fluor 488 (white) was injected into the brain parenchyma of surface fish. Microglial lysosomes were labeled with LysoTracker Red (magenta) and imaged by confocal time-lapse microscopy. Images were acquired every 1.5 min and are displayed at 3 frames per second.

