## Supplementary figures and images for "Evolved differences in microglial cell biology between surface and cave populations of *Astyanax mexicanus*"

### Figure S1

**Supplementary figure 1**

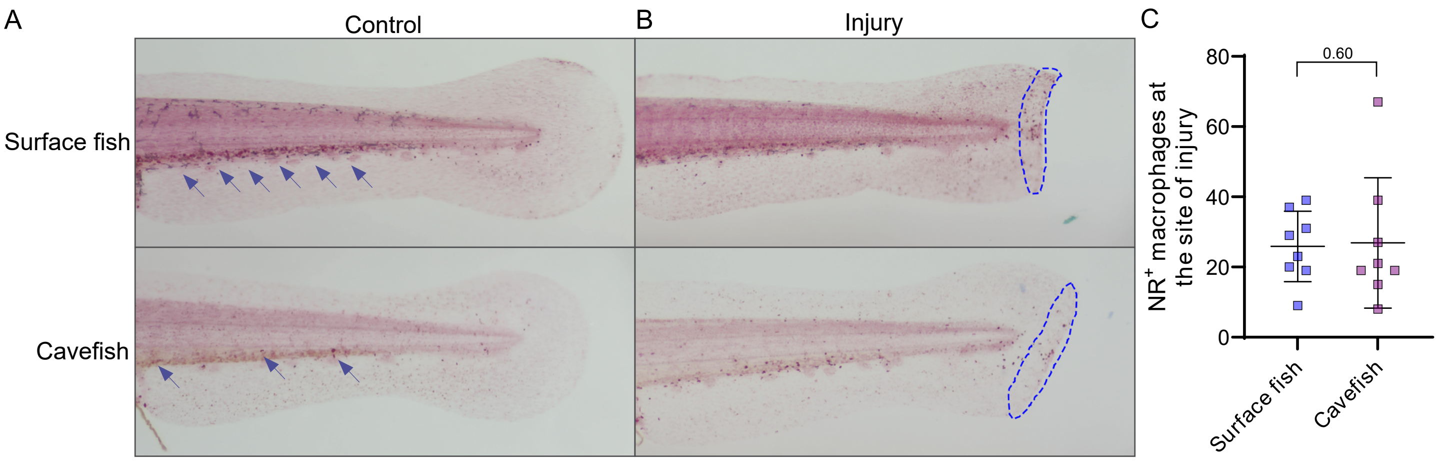

### Figure S2

**Supplementary figure 2**

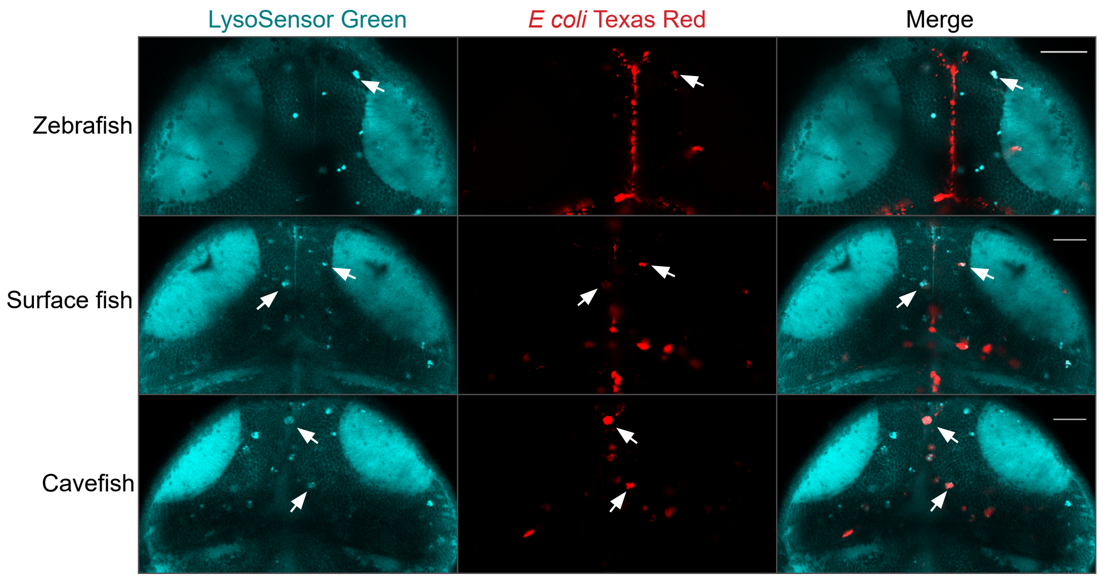
